# A Distinct Alternative mRNA Splicing Profile Identifies the Oncogenic CD44 Transcript Variant 3 in KMT2A-Rearranged Pediatric T-cell Acute Lymphoblastic Leukemia Cells

**DOI:** 10.1101/2024.09.05.611385

**Authors:** Amanda Ramilo Amor, Sabina Enlund, Indranil Sinha, Qingfei Jiang, Ola Hermanson, Anna Nilsson, Shahrzad Shirazi Fard, Frida Holm

## Abstract

T-cell acute lymphoblastic leukemia (T-ALL), which constitutes of 10-15% of all pediatric ALL cases, is known for its complex pathology due to pervasive genetic and chromosomal abnormalities. Although most children are successfully cured, chromosomal rearrangements involving the KMT2A (KMT2A) gene is considered a poor prognostic factor. In a cohort of 171 pediatric T-ALL samples we have studied differences in gene and splice variant patterns in KMT2A rearranged (KMT2A-r) T-ALL compared to KMT2A negative (KMT2A-wt) T-ALL samples. Our results have identified a distinct gene expression and splice variant expression pattern in pediatric KMT2A-r patient samples including significant expression of splicing regulatory markers ESRP1 and MBNL3. Additionally, the pro-survival long transcript variant of BCL2 were upregulated in KMT2A-r compared to KMT2A-wt T-ALL samples. Lastly, increased levels of activating methylation in the promoter region of CD44 were identified followed by an upregulation of the oncogenic transcript variant CD44v3 in KMT2A-r T-ALL. Together this suggests that CD44v3 could play a potential role as gene expression-based risk stratification of KMT2A-r rearranged T-ALL and could possibly serve as a therapeutic target using splicing modulators.

## 1. INTRODUCTION

The development of novel therapies for hematological malignancies characterized by unfit stem cell activity represents an urgent unmet medical need, particularly in pediatric malignancies with high relapse rates and potential fatal late effects as a direct results of aggressive cancer therapies. Several hematological malignancies, such as T-cell-acute lymphoblastic leukemia (T-ALL) is often enriched with leukemia initiating cells (LICs), which exhibit enhanced survival and self-renewal capacities^1-3^. However, the mechanism by which LICs gain survival and self-renewal advantages are largely unknown. Uncovering these mechanisms is critical for identifying targeted approaches to eliminate LICs and developing curative therapies.

Advances in cancer biology has revealed that RNA modifications such as alternative splicing are one of the main drivers for cancer initiation and relapse^4^. Alternatively spliced genes generate mature messenger RNA (mRNA) which depending on the exon sequence can code for different transcript variants, or isoforms. Transcript variants generated from the same gene can be translated into different proteins with altered functions. Thus, it is not surprising that disruption in the alternative splicing program can promote the development of disease.

Rearrangement of mixed-lineage leukemia (MLL), sometimes referred to as lysine (K)-specific methyltransferase 2A (KMT2A) is one of the major driver mutations in acute leukemia. In general, KMT2A rearrangement (KMT2A-r) is associated with poorer prognosis in both acute lymphoblastic leukemia (ALL) and acute myeloid leukemia (AML)^5^. However, the regulatory network of genes involved in. KMT2A-r driven T-ALL is unknown. Here we set out to map genes associated to KMT2A-r and to identify possible transcript variants of known genes involved in cancer stem cell maintenance. One possible marker is transcript variants of the adhesion molecule CD44, which previously have been shown to be overexpressed in several cancer types and is considered a common marker for cancer stem cells^6,7^. To date, the role of CD44 in high-risk T-ALL has not been fully elucidated. This is important since current chemotherapy approaches target pathways essential for general cell survival but not the specific driver mutations. Additionally, current therapeutic approaches only hit leukemia bulk but not the LICs, leaving possibilities for relapse from the malignant clones surviving initial treatment protocols. This could potentially contribute to the high relapse rate and poor long-term survival rate of KMT2A-r leukemia patients.

## 2. METHODS

### 2.1. RNA sequencing analysis

For RNA-sequencing analysis, 161 KMT2A-wt T-ALL and 10 KMT2A-r T-ALL samples were selected; samples are not longitudinal. The RNA-sequencing dataset (.BAM) files sourced from T-ALL patients were acquired via the GDC Data Portal (https://portal.gdc.cancer.gov/projects/TARGET-ALL-P2). For the whole gene expression data, we used the GDC mRNA Analysis Pipeline output, which was downloaded from the GDC data portal. Fragments per Kilobase of transcript per Million mapped reads (FPKM) values for each sample were downloaded and merged for further analysis. Downloading and generation of raw count values for transcript variant expression was established in SeqMonk Mapped Sequence Data Analyzer tool (version 1.47.1) according to the pipeline used in Enlund et al^8^. Data is presented as Frames per Kilo-base of exon per Million mapped reads (FPKM) values normalized in R (version 4.3.1) using the transformation guideline provided by the GDC bioinformatics pipeline. Principal Component Analysis (PCA) and heatmaps were created using Qlucore Omics Explorer (version 3.9). Gene-set Enrichment Analysis (GSEA) and over-representative analysis (ORA) was generated through R (version 4.3.1), using packages clusterProfiler (version 4.8.3) and enrichplot (version 1.20.3). None-coding genes such as novel transcripts, long intergeneic non-protein coding (LINC), novel proteins and pseudogenes where removed from the analysis.

### 2.2. Cell lines

T-ALL cell lines Jurkat and CCL-119 (ATCC) were cultured in suspension in RPMI with 10% FBS (Sigma-Aldrich). T-ALL cell line MOLT16 (Leibniz Institute DSMZ) was cultured in suspension in IMDM (Gibco) with 10% FBS. KMT2A-r cell line KARPAS-45 was cultured in suspension using RPMI supplemented with 2mM glutamine and 20% FBS (Sigma-Aldrich). KMT2A-r cell line SUP-T13 was cultured in suspension in RPMI (ATCC modification) (Gibco) with 10% FBS and penicillin 100 U/ml and streptomycin 100 μg/ml.

### 2.3. Primary patient samples

Mononuclear cells from primary T-ALL samples were obtained after informed consent and in accordance to the **Ethical permission: dnr 2021-02718**. Samples underwent positive CD34 selection using EasySep Human CD34 positive Selection kit II (Stem Cell Technologies).

### 2.4. RNA preparation and qRT-PCR analysis

RNA preparation and qRT-PCR analysis were performed as described in Enlund et al^8^. Splicing specific primer sequences for CD44v3 and the BCL2 family members can be found in Holm et al^6^.

### 2.5. Chromatin Immunoprecipitation

Chromatin ImmunpPrecipiation (ChIP) was performed using the Diagenode True Microchip kit (Diagenode), following the manufacturer’s protocol. Data is presented as fold enrichment.

### 2.6. Statistical analysis

Statistical analyses were performed by multiple and standard unpaired t-test when comparing two groups. Significance was set to p≤0.05 (*p ≤ 0.05, **p<0.01, ***p<0.001, ****p<0.0001). Results are presented as the mean ± SEM.

## 3. RESULTS and DISCUSSION

### 3.1 KMT2A-r T-ALL patients are distinguished by dysregulation of genes that play a protumor role through transcriptional misregulation

To determine alterations in gene expression in pediatric KMT2A-r T-ALL, a total of 171 bone marrow derived samples were included (Figure 1A, Supplemental Figure S1A). Samples categorized as KMT2A-wt were negative for any other known mutation. Whole genome RNA-sequencing analysis revealed dysregulated genes uniquely differentially expressed, distinguishing patients with KMT2A-r from KMT2A-wt T-ALL patients (Figure 1B). 15% of the genes were significantly differentially expressed in KMT2A-r T-ALL patient samples compared to KMT2A-wt T-ALL, most of which (88%) were upregulated (Supplemental Figure S1B). The differential expression patterns of KMT2A-r T-ALL patients were further supported by PCA, where KMT2A-r patients and KMT2A-wt patients were separated into distinct clusters (Figure 1C). Interestingly, patient samples from the KMT2A-r group reveled an upregulation of MIR1915 (Figure 1D). MIR1915 has been described to play a role in many malignancies such as gastric cancer^9^, colorectal carcinoma^10^ and breast cancer^11^. MIR1915 has previously been shown to play a key role in drug resistance by targeting apoptotic regulatory genes in the B-cell lymphoma 2 (BCL-2) family^12,13^. Additionally, oncogenes TCF4 and SKIDA1 (Figure 1D) where highly upregulated in KMT2A-r T-ALL patient samples. To further elucidate the role of the differential gene expression profile, GSEA was performed to identify dysregulated pathways and genes in KMT2A-r T-ALL patients compared to KMT2A-wt T-ALL patients. Interestingly, the GSEA revealed enrichment of transcriptional misregulation in cancer and ORA revealed several genes in the pathway significantly expressed in KMT2Ar T-ALL compared to KMT2A-wt patients (Figure 1E-F). Taken together, rearrangement in KMT2A correlates with expression alterations in genes important for malignancy and drug resistance.

**Figure 1.**
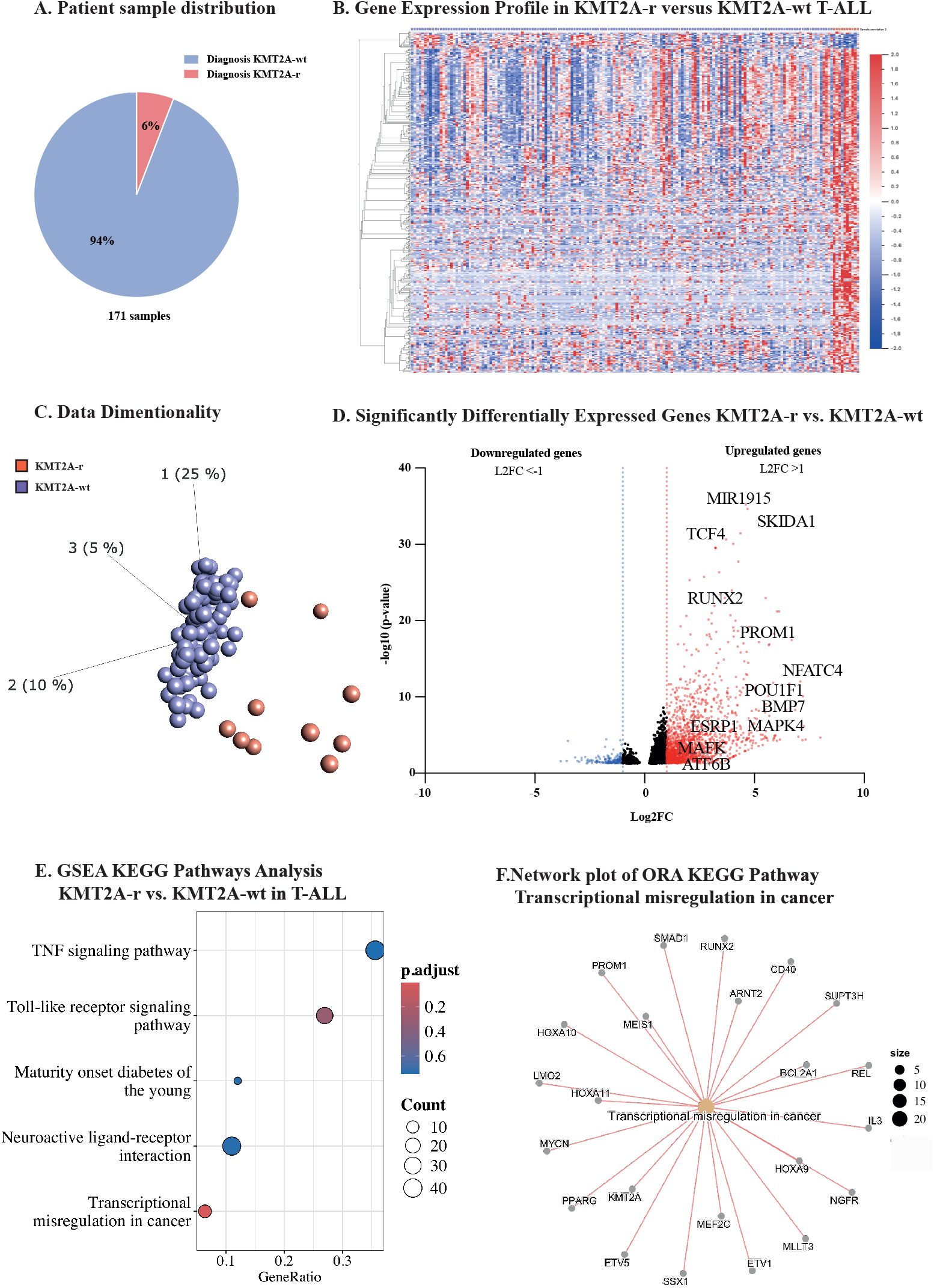
Gene signature of KMT2A-rearranged T-ALL and KMT2A-wt T-ALL. **A)** Schematic overview of included T-ALL patients from the TARGET database. Samples were distributed as 5.85% KMT2A-r T-ALL (n=10) and 94.15% KMT2A-wt T-ALL (n=161) and include only bone marrow samples. **B)** Heat map of top 400 significantly differentially expressed genes in KMT2A-r T-ALL compared to KMT2A-wt samples. Significance calculated by unpaired student’s t-test (p<0.05). **C)** PCA plot based on gene expression of diagnosis samples classified as KMT2A-wt or KMT2A-r generated using in Qlucore Omics Explorer (version 3.9). Significance calculated unpaired students t-test (p<0.05, SD<0.25). **D)** Volcano plot of top significantly differentially expressed genes in KMT2A-r compared to KMT2A-wt samples, displayed as L2FC calculated from normalized FPKM values. Significance calculated by unpaired student’s t-test (p<0.05). **E)** GSEA KEGG Pathway Analysis of all genes in KMT2A-r T-ALL compared to KMT2A-wt T-ALL, rank calculated by negative log10 p-value and signed L2FC. For Transcriptional misregulation: NES=1,33, p-value=2,6E-05, p.adjust=0,0076. Significance displayed as adjusted p-value. **F)** Network plot of the pathway transcriptional misregulation in cancer created through ORA using KEGG pathways of differentially expressed genes (p<0.05, L2F1>1) in KMT2A-r T-ALL compared to KMT2A-wt T-ALL.

### 3.2 Malignant upregulation of CD44 transcript variant 3 (CD44v3) in KMT2A-r T-ALL patient samples

RNAs, including microRNAs (miRNA), are known for their role in cancer progression by regulating tumor cell growth, migration and invasion, as well as apoptosis. Although a highly significant upregulation of MIR1915 was identified (Figure 2A, left panel) no dysregulation of the BCL2 gene (Figure 2A, right panel) or other family members (Supplemental Figure S1C) were identified. Interestingly, we identified an upregulation of Epithelial-Splicing-Regulatory-Protein 1 (ESRP1) in KMT2A-r T-ALL patients samples (Figure 2B left). ESRP1 is an RNA-binding protein essential for mammalian development by regulating splicing during epithelial to mesenchymal transmission (EMT)^14^. Through alternative mRNA splicing, different transcript variants from the same gene can be generated, which can be translated into different proteins with altered functions. In the light of this, we investigated and identified a distinct transcript variant landscape in KMT2A-r samples (Supplemental Figure S1D-E). Splicing of apoptosis regulatory genes BCL2, BCL2L1 (BCLX), and MCL1 produces two distinct isoforms, where the long transcript variants of promote cell survival, while the short transcript variants are pro-apoptotic^15,16^. Expression of the long pro-survival transcript variant BCL2-L was significantly upregulated in KMT2A-r T-ALL compared to KMT2A-wt T-ALL (Figure 2B right). Hence, pro-survival transcript variant switching of BCL2 may have clinical utility in predicting malignant transformation promoting a phenotype with pro-survival characteristics in KMT2A-r T-ALL patients. Contrarily, in KMT2A-wt T-ALL the long transcript variants of both BCLX and MCL1 is significantly upregulated (Supplemental Figure S1F). ORA of gene ontologies identified several biological processes dysregulated in KMT2A-r T-ALL. Among these we identified cell-cell adhesion via plasma-membrane adhesion molecules and homophilic cell adhesion via plasma membrane adhesion molecules (Figure 2C). One specific cell surface adhesion molecule, CD44, has been widely implicated as a cancer stem cell marker^17,18^. Several positive regulators are significantly upregulated in KMT2A-r T-ALL patient samples, such as T-cell factor 4 (TCF4), a Wnt/b-catenin signaling pathway activator expressed in multiple cancers and Mitogen-activated protein kinase 3 (MAPK3)^19,20^ (Figure 2E). In contrast, none of the negative regulators, TP53, KLF4 or FOXP3, where differentially expressed (Supplemental Figure S1G). The upregulated splicing regulator ESRP1 have previously been identified to regulate the adhesion molecule CD44 transcript variant expression^14^ (Figure 2F). Transcript variants of the adhesion molecule CD44, specifically CD44 transcript variant 3 (CD44v3) have previously been shown to be overexpressed in hematological malignancies^6^, and is considered a common marker for cancer stem cells. Interestingly, in T-ALL cell lines positive for KMT2A rearrangement we could show increased levels of activating methylation in the promoter region of CD44 compared to cell lines negative for the KMT2A rearrangement using ChIP coupled with RT-qPCR (Figure 2G). Although CD44 is not significantly upregulated in KMT2A-r T-ALL (Figure 2 H, upper left panel), the negative splicing regulator of CD44, muscleblind like splicing regulator 3 (MBNL3), is significantly downregulated (Figure 2H, upper right panel). With two significantly differentially expressed regulators of CD44 alternative splicing (ESRP1 and MBNL3), the significant upregulation of CD44v3 in KMT2A-r T-ALL patient samples (Figure 2H lower left panel, Supplemental Figure S1H) and in CD34+ selected stem cells from pediatric primary patients with T-ALL (Figure 2H lower right panel) should be considered a potential cancer stem cell biomarker for KMT2A-r T-ALL.

**Figure 2.**
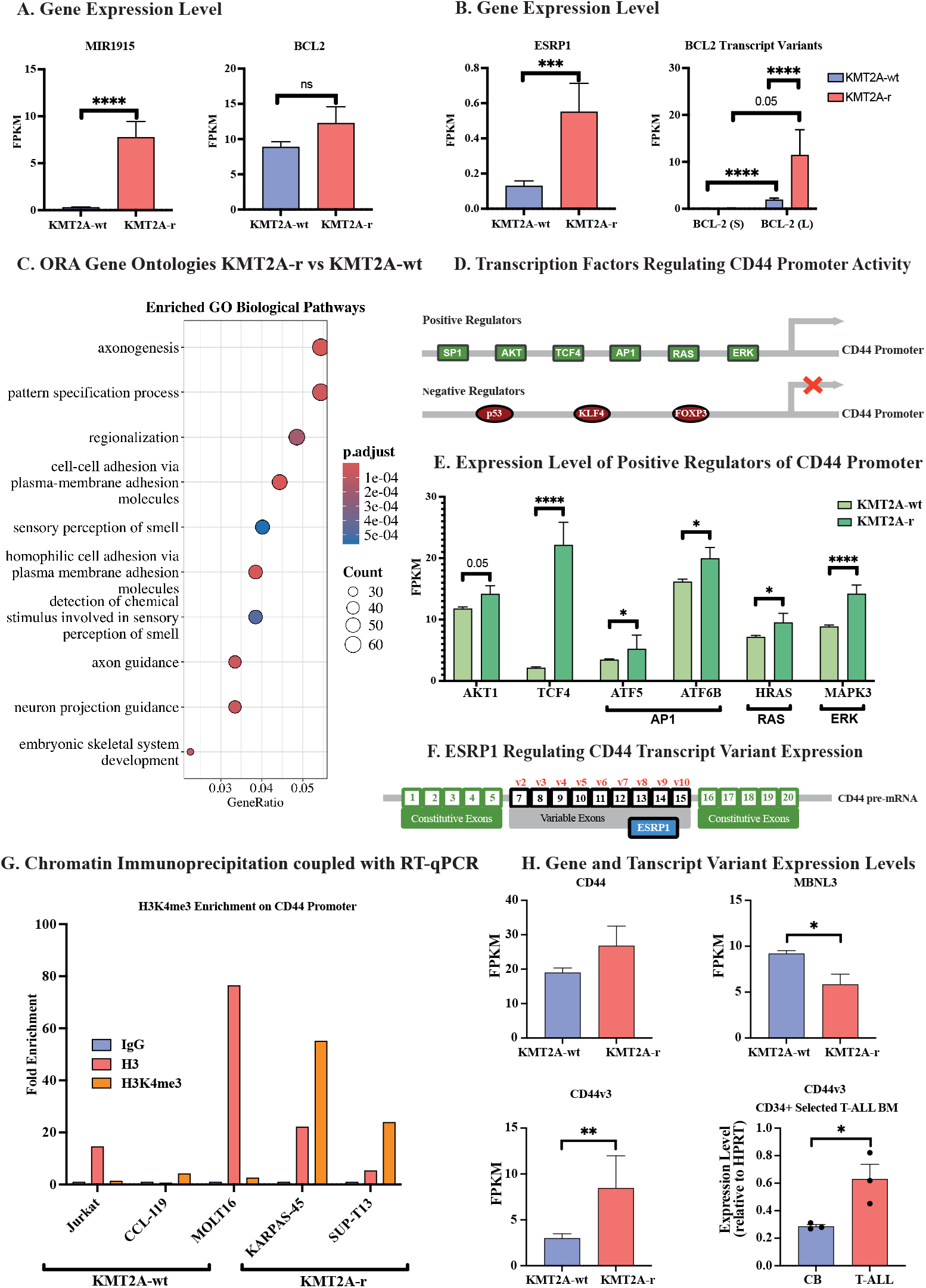
Distinct alternative mRNA splicing pattern resulting in CD44v3 upregulation in KMT2A-r T-ALL. **A)** Gene expression levels displayed as FPKM of MIR1915 and BCL2 in pediatric KMT2A-r T-ALL compared to KMT2A-wt T-ALL. Significance calculated by unpaired students t-test (*p < 0.05). **B)** Gene expression levels displayed as FPKM of ESRP1 (left) and transcript variant expression level of BCL2 transcript variant in pediatric KMT2A-r T-ALL compared to KMT2A-wt T-ALL. Significance calculated by unpaired students t-test (*p < 0.05). **C)** ORA of differentially expressed genes (p<0.05, L2F1>1) showing enrichment of the gene ontology biological processes in KMT2A-r T-ALL compared to KMT2A-wt T-ALL. **D)** Schematic visualization of the positive and negative regulators of CD44 promoter activity modified from^21^. **E)** Gene expression levels displayed as FPKM of the positive regulators of CD44 promoter activity in pediatric KMT2A-r T-ALL compared to KMT2A-wt T-ALL. Significance calculated by unpaired students t-test (*p < 0.05). **F)** Schematic visualization of ESRP1s role in regulating CD44 transcript variant expression, modified from^21^. **G)** Chromatin Immunoprecipitation coupled with RT-qPCR showing increased expression of H3K4me3 on the CD44 promoter region in T-ALL cell lines. Results displayed as fold enrichment (compared to IgG/Mock). **H)** Gene expression levels in pediatric KMT2A-r T-ALL compared to KMT2A-wt T-ALL displayed as FPKM of CD44 (top left panel), MBNL3 (top right panel) and CD44v3 (bottom left panel). Additionally, CD44v3 transcript variant expression in CD34+ selected primary T-ALL patient samples by RT-qPCR, normalized to HPRT (bottom right panel). Significance calculated by unpaired students t-test (*p < 0.05).

## 4. ACKNOWLEDGEMENT

This work was supported by the Swedish Childhood Cancer Foundation (Barncancerfonden; F.H), Åke Wiberg (F.H.) and Märta och Gunnar V. Philipson Foundation (A.N and F.H.). We would like to thank Dr. Rozbeh Jafari at Karolinska Institutet and Professor Jonathan Pollack at Stanford University for kindly sharing T-ALL cell lines with us.

The TARGET Initiative (RNA-sequencing data-set): The results published here are in whole or part based upon data generated by the Therapeutically Applicable Research to Generate Effective Treatments (TARGET) initiative, phs000218, managed by the NCI. The data used for this analysis are available at https://portal.gdc.cancer.gov/projects/TARGET-ALL-P2.

Information about TARGET can be found at http://ocg.cancer.gov/programs/target. Part of the data handling was enabled by resources at project number SNIC-2021/22-48 provided by the Swedish National Infrastructure for Computing (SNIC) partially funded by the Swedish Research Council through grant agreement no. 2018-05973.

## 5. AUTHORSHIP STATEMENT

Amanda Ramilo Amor, Sabina Enlund and Indranil Sinha collected and analyzed the data. Sabina Enlund and Frida Holm conceptualized the project, analyzed data, and wrote the paper. Anna Nilsson, Ola Hermanson and Shahrzad Shirazi Fard contributed with scientific expertise and were involved in writing and reviewing the paper.

## FIGURE LEGEND

**Supplemental figure 1**.

**A)** Cohort characterization of included samples.

**B)** Schematic overview of the distribution of genes (left panel) between KMT2A-r T-ALL (n=10) and KMT2A-wt T-ALL (n=161) and if expression is up- or downregulated (right panel).

**C)** Gene expression levels displayed as FPKM of BCLX and MCL1 in pediatric KMT2A-r T-ALL compared to KMT2A-wt T-ALL. Significance calculated by unpaired students t-test (*p < 0.05).

**D)** Heat map of top 400 significantly differentially expressed transcript variants in KMT2A-r T-ALL compared to KMT2A-wt samples. Significance calculated by unpaired student’s t-test (p<0.05).

**E)** PCA plot of transcript variants of diagnosis samples classified as KMT2A-wt or KMT2A-r generated using in Qlucore Omics Explorer (version 3.9). Significance calculated by multi group ANOVA (p<0.05, SD<0.25).

**F)** Splice variant expression level of BCLX and MCL1 transcript variant in pediatric KMT2A-r T-ALL compared to KMT2A-wt T-ALL. Significance calculated by unpaired students t-test (*p < 0.05).

**G)** Gene expression levels displayed as TPM of the negative regulators of CD44 promoter activity (TP53, KLF4 and FOXP3) in pediatric KMT2A-r T-ALL compared to KMT2A-wt T-ALL. Significance calculated by unpaired students t-test (*p < 0.05).

**H)** Splice variant expression level of protein coding CD44 transcript variants in pediatric KMT2A-r T-ALL compared to KMT2A-wt T-ALL. Significance calculated by unpaired students t-test (*p < 0.05).

