## Supplemental figure 1 for "A Distinct Alternative mRNA Splicing Profile Identifies the Oncogenic CD44 Transcript Variant 3 in KMT2A-Rearranged Pediatric T-cell Acute Lymphoblastic Leukemia Cells"

### A. Cohort Characterization T-ALL (TARGET)

| Variables | Acute leukemia T-ALL | MLL Status KMT2A-wt | KMT2A-r |
|---|---|---|---|
| ***Age group** | | | |
| Pediatric (1-15 years old) | 155 | 125 | 8 |
| Older (15-30 years old) | 16 | 36 | 2 |
| **Sex** | | | |
| Female | 43 | 40 | 3 |
| Male | 128 | 121 | 7 |
| **Sample type** | | | |
| Bone Marrow | 171 | 161 | 10 |
| Peripheral Blood | 0 | 0 | 0 |
| **MLL status** | | | |
| KMT2A-wt | 161 | - | - |
| KMT2A-r | 10 | - | - |
| **Total** | 171 | 161 | 10 |

### B. Differentially Expressed Genes

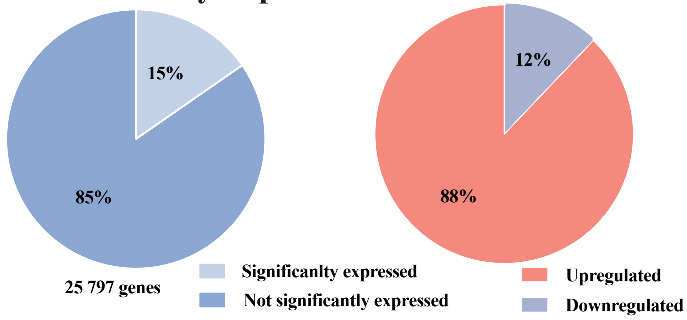


### C. BCL2 Family Gene Expression

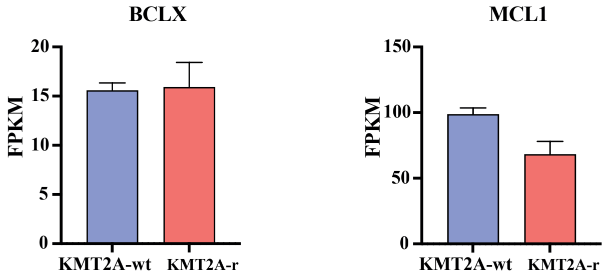


### D. Transcript Variant Signature KMT2A-r vs KMT2A-wt

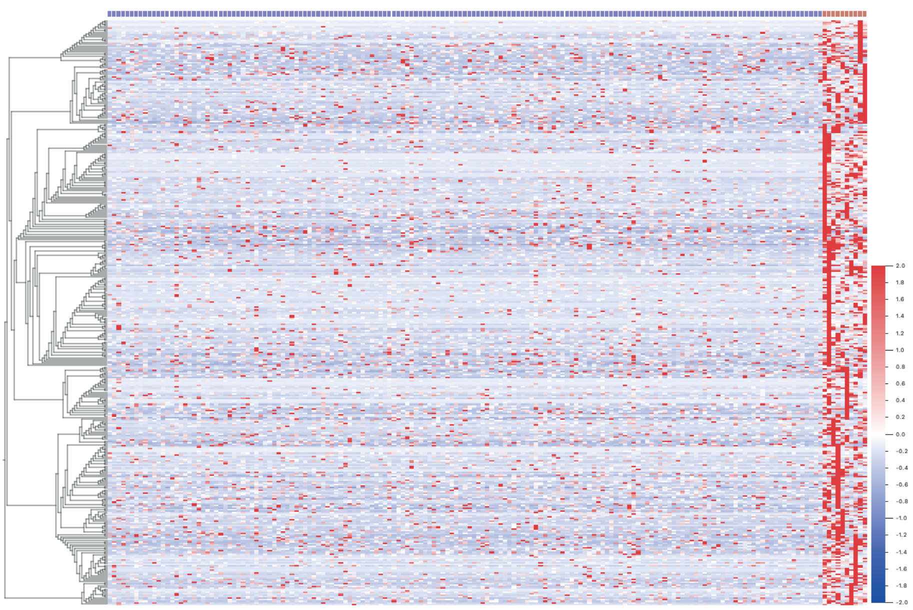

### E. PCA Plot of Samples by KMT2A status

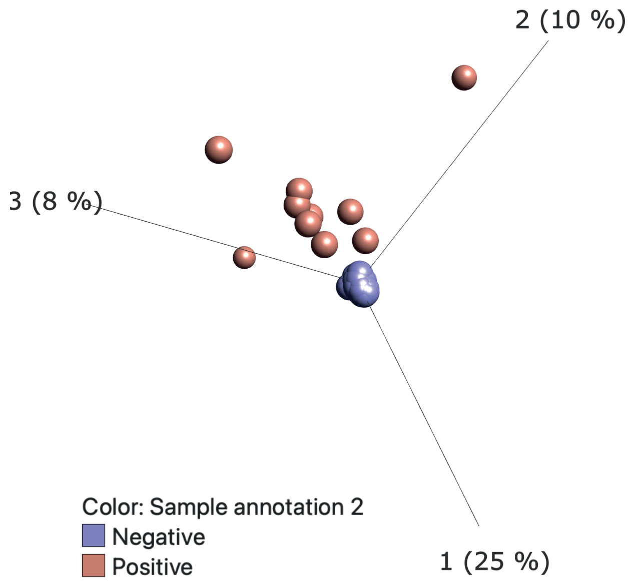


### F. BCL2 Family Transcript Variant Expression

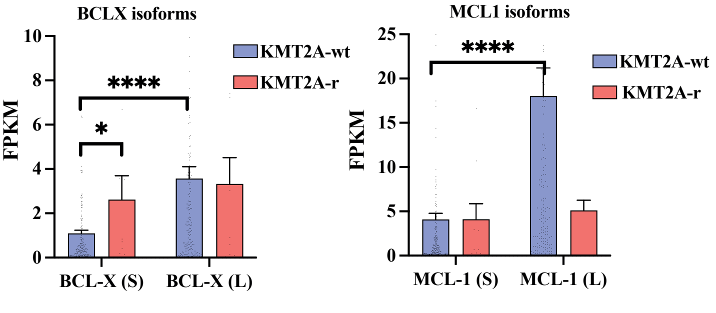


### G. Expression of Negative Regulators of CD44 Promoter

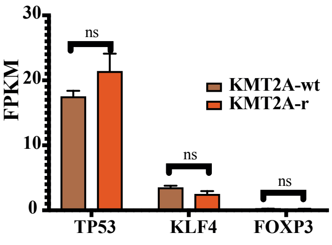


### H. Protein Coding CD44 Transcript Variant Expression

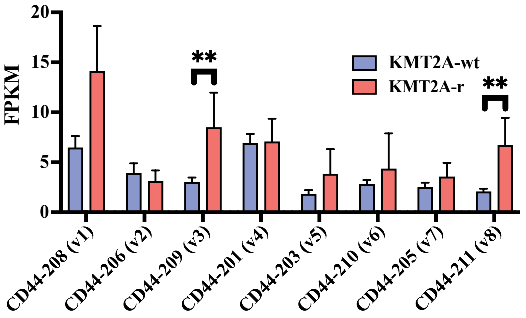
